## Supplementary material for "Protocol and reagents for pseudotyping lentiviral particles with SARS-CoV-2 Spike protein for neutralization assays": File S1 (plasmid maps in Genbank format): README.md

# Plasmid maps
Maps of plasmids for SARS-CoV-2 lentiviral pseudotyping.
The links go to Genbank plasmid maps that can be opened in programs such as [ApE](https://jorgensen.biology.utah.edu/wayned/ape/) or [Benchling](https://www.benchling.com/).

 - [HDM_IDTSpike_fixK](2710_HDM_IDTSpike_fixK.gb): plasmid expressing the Spike protein from SARS-CoV-2 with the sequence codon optimized (#2710 in Bloom lab plasmid log).

 - [HDM_IDTSpike_fixK_HA_Tail](2707_HDM_IDTSpike_fixK_HA_Tail.gb): plasmid expressing the Spike protein from SARS-CoV-2 with the seuqence codon optimized and the cytoplasmic tail replaced by that from influenza HA (#2707 in Bloom lab plasmid log).

 - [HDM-nCoV-Spike-IDTopt-ALAYT](2702_HDM-nCoV-Spike-IDTopt-ALAYT.gb): plasmid expressing the Spike protein from SARS-CoV-2 with the sequence codon-optimized by IDT and two mutations in the last five amino-acids (so that the C-terminal sequence is ALAYT). This is plasmid #2703 in Bloom lab plasmid log.

 - [pHAGE2_CMV_ZsGreen_W](25_pHAGE2_CMV_ZsGreen_W.gb): lentiviral backbone expressing ZsGreen under a CMV promoter (#25 in Bloom lab plasmid log).

 - [pHAGE-CMV-Luc2-IRES-ZsGreen-W](2728_pHAGE-CMV-Luc2-IRES-ZsGreen-W-1270.gb): lentiviral backbone expressing Luciferase followed by an IRES-ZsGreen under a CMV promoter (#2728 in Bloom lab plasmid log).

 - [pHAGE2_EF1aInt_ACE2_WT](2653_pHAGE2_EF1aInt_ACE2_WT.gb): lentiviral backbone expressing human ACE2 under a EF1a promoter containing an intron (#2653 in Bloom lab plasmid log).

 - [HDM_Hgpm2](26_HDM_Hgpm2.gb): lentiviral helper plasmid expressing HIV Gag-Pol under a CMV promoter (#26 in Bloom lab plasmid log).

 - [HDM_tat1b](27_HDM_tat1b.gb): lentiviral plasmid expressing HIV Tat under a CMV promoter (#27 in Bloom lab plasmid log).

 - [pRC_CMV_Rev1b](28_pRC_CMV_Rev1b.gb): lentiviral plasmid expressing HIV Rev under a CMV promoter (#28 in Bloom lab plasmid log).
